# Intrinsic Reproductive Isolation among Ecologically Divergent *Anolis* Lizards

**DOI:** 10.1101/2025.08.14.670316

**Authors:** Anthony J. Geneva, Richard E. Glor

## Abstract

The evolution of reproductive isolation lies at the core of our modern concept of species. Despite this central importance, study of the evolution of reproductive isolation is largely limited to a set of laboratory amenable species. When shallowly divergent populations are associated with different habitats, extrinsic isolation is often assumed to play a major role. Because this type of isolation is environmentally mediated it can be ephemeral, for instance, if habitats change in response to climate. In contrast, intrinsic isolation is heritable and therefore loss of this type of isolation requires genetic changes in one or both populations. We performed a hybridization and backcross experiment to test for and characterize intrinsic reproductive isolation between two recently diverged, ecologically differentiated populations of *Anolis* lizards. We find evidence of substantial intrinsic isolation that appears to operate at the postmating, prezygotic phase. Our findings suggest that intrinsic isolation may play a substantial role in the maintenance of shallow, yet ecologically divergent, lineages.

## 1 Introduction

The gold standard for species diagnosis is the existence of irreversible, genetically-based barriers that prevent reproduction under any conditions (Dobzhansky 1935; Mayr 1963). Such intrinsic reproductive isolation has been reported across taxa ranging from plants to rodents and was, until relatively recently, the basis of most genetical models of speciation (Coyne and Orr 2004; Fishman and Willis 2001; Fitzpatrick 2008; Moyle and Nakazato 2009; Turelli et al. 2001). The degree to which most animal species are intrinsically isolated, however, is completely unknown; even the strongest proponents of the biological species concept – which defines species as reproductively isolated populations – tend to diagnose and delimit species using proxies for reproductive isolation such as phenotypic or genetic differentiation (e.g., Mayr 1992; Whittemore 1993). Direct assessment of reproductive isolation is scarcely mentioned in recent reviews of methods for delimiting species in nature (Camargo and Sites 2013; Sites and Marshall 2003). Because it is so difficult to investigate in nature, much of the information we do have about intrinsic reproductive isolation is derived from laboratory model organisms or domesticated species.

Intrinsic reproductive isolation is not necessary for speciation; indeed, a growing number of studies emphasize the importance of extrinsic reproductive isolation, which occurs when species capable of successful reproduction do not interbreed in nature due to non-overlapping distributions, ecological differences, or other context dependant isolating mechanisms (Coyne and Orr 2004). Extrinsic isolation is thought to be particularly important between relatively young species where rapid divergence of sexually selected or adaptive traits associated with reproductive isolation (e.g., ecological speciation) may outpace more gradual accumulation of genetic changes often expected of intrinsic isolation (Hendry et al. 2007; Nosil 2012; Schluter 2009; Sobel et al. 2010). Although intrinsic and extrinsic isolation are not mutually exclusive, and often appear to act in concert (Nosil et al. 2005; Ramsey et al. 2003), recent studies of species collapse following environmental changes suggest that reproductive isolation between some species may be largely or even entirely extrinsic (reviewed in Seehausen et al. 2008). For example, decreased visibility due to human activity can interfere with visual mating preferences in Lake Victoria cichlid fish, resulting in the breakdown of assortative mating and collapse of some species into hybrid swarms (Seehausen et al. 1997).

We focus our attention here on assessing intrinsic isolation in a pair of relatively recently diverged anole lizards. Anoles are an ideal system for investigating the role of intrinsic isolation early in the speciation process because divergence of sexual signals and adaptive traits in this taxon is expected to result in extrinsic reproductive isolation (Losos 2009; Schuluter 2000; Williams 1972). We mitigate the influence of extrinsic barriers by using a common garden to test whether intrinsic isolation has evolved between young, ecologically differentiated lineages. Evidence of hybridization among anole species is extremely rare in nature, suggesting the existence of multiple persistent and strong isolating mechanisms between most species (Losos 2009; Ng et al. 2017). Despite this, preliminary evidence for reproductive isolating barriers exists for only two pairs of anole species, both involving a type of intrinsic barrier — chromosomal abnormalities in hybrid progeny (Gorman et al. 1971; Webster 1977).

We test the hypothesis that intrinsic isolation is present between anole taxa by quantifying a suite of reproductive variables indicative of intrinsic isolation (viable and inviable egg production, egg incubation success, hatchling survival, and hatchling sex ratio) and testing if these variables support the prediction of significantly reduced fitness for hybrid crosses. By quantifying the strength of intrinsic reproductive isolation we are also able to compare the degree of intrinsic isolation between this pair to that observed between other well-studied species sampled from across the tree of life.

In addition to assessing the presence and strength of intrinsic isolation, we also distinguish among three possible types of intrinsic isolation that act at different stages of the reproductive process. Distinguishing among these types of intrinsic isolation is important because they result from different evolutionary mechanisms affecting distinct groups of genes and biological processes. The earliest acting type of intrinsic isolation occurs when individuals are unable to recognize heterospecifics as potential mates. Such intrinsic premating isolation is important in frogs, birds, insects and other animals where speciation results from divergence of signals important to species recognition and sexual selection (Grant and Grant 1997; Littlejohn 1965; Smadja and Butlin 2009). Intrinsic premating isolation in anoles may involve divergence in dewlap color and pattern, or divergence in the complicated stereotypical display repertoires during which dewlaps are extended and retracted (Leal and Fleishman 2004; Losos 1985; Ng et al. 2012; Williams and Rand 1977). Intrinsic premating isolation predicts reduced fitness of hybrid crosses due to reduced mating frequency. We test this prediction by determining if hybrid crosses produce significantly fewer eggs overall (including both viable and inviable) than pure crosses during the first generation of our experiment. Virgin female anoles only occasionally produce inviable eggs, whereas mated females regularly produce two to four single egg clutches per month (Andrews and Rand 1974).

A second type of intrinsic isolation occurs after mating, but prior to fertilization (Birkhead et al. 1994). Such postmating, prezygotic (PMPZ) isolation generally results from tight coevolution between male and female reproductive systems (Eady 2001). Examples include *Drosophila* sperm that are transferred but cannot be properly stored in the reproductive tract of heterospecific females and broadcast spawning abalone where sperm lysins are unable to dissolve the vitelline envelope of heterospecific eggs (Price et al. 2001; Vacquier and Lee 1993). PMPZ isolation is now recognized as an important form of reproductive isolation across a diverse range of arthropods, bird, and plant species (Birkhead and Brillard 2007; Larson et al. 2012; Scopece et al. 2007; Sweigart 2010; Wade et al. 1994). The fact that reproductive proteins are among the fastest evolving proteins across the tree of life further suggests that this type of isolation may be more common and quick to evolve than previously expected (Swanson and Vacquier 2002). Such rapid evolution may be due to coevolutionary arms races driven by sexual conflict (Birkhead and Brillard 2007; Eady 2001; Gavrilets 2000; Parker and Partridge 1998; Swanson and Vacquier 2002). Although PMPZ has not been studied previously in anoles, they do exhibit at least three forms of sexual conflict. First, sperm competition may be common in female anoles have been frequently found to mate with more than one male (Calsbeek et al. 2007; Johnson 2007; Kamath and Losos 2017; Passek 2002). Second, post-copulatory cryptic sperm choice has been reported in *Anolis sagrei* (Calsbeek and Bonneaud 2008). Third, *A. carolinensis* females experience coition-induced inhibition of sexual receptivity, a similar phenotype in flies causes sexual conflict and may have led to antagonistic sexual co-evolution (Chapman 2001; Crews 1973; Knowles and Markow 2001; Sweigart 2010; Wigby and Chapman 2005).

Although PMPZ isolation is difficult to observe, particularly in internally fertilized species (where it occurs entirely in the reproductive tract of females), it may be distinguished from premating and postzygotic isolation in anoles and other egg laying squamates (Corl et al. 2012). Female anoles that have not mated tend not to produce any eggs, whereas post-mating fertilization failure is expected to result in uncalcified eggs that are easily distinguished from the calcified shells typical of fertilized eggs (Corl et al. 2012; Olsson et al. 1994; Olsson and Shine 1997; Sinervo and Licht 1991). PMPZ isolation may be responsible for the increased proportion of uncalcified eggs produced in crosses between populations of the side-blotched lizards (Corl et al. 2012). We test for PMPZ isolation in anoles by asking if fewer viable eggs are produced by first generation hybrid crosses than pure crosses.

The third and final type of intrinsic reproductive isolation involves hybrid sterility or inviability (reviewed in Coyne and Orr 2004). The best known example of this type of intrinsic isolation involves the production of sterile mule offspring from crosses between female horses and male donkeys. This intrinsic postzygotic isolation is thought to evolve as the result of Dobzhansky-Muller incompatibilities — the accumulation of genetic differences in separate lineages which result in reduced fitness of hybrids due to negative epistatic interactions between genotypic combinations that do not occur in either parental population (Orr 1995). This type of incompatibility relies on the accumulation of genetic differences and can evolve rapidly through changes in genes associated with adaptive evolution or intragenomic conflict, or more slowly through the stochastic accumulation of neutral substitutions throughout the genome over time (Orr and Turelli 2001; Presgraves 2010; Turelli et al. 2001). For example, postzygotic hybrid male sterility between *Drosophila simulans* and *D. melanogaster* is the result of positive natural selection on two genes, one in each species (Barbash et al. 2003; Brideau et al. 2006). In anoles, two small cytological studies of field collected hybrids found meiotic failure in hybrid males resulting in intrinsic postzygotic isolation in the form of hybrid sterility (Gorman et al. 1971; Webster 1974, 1977). Intrinsic postzygotic isolation predicts reduced viability or fertility in the offspring of hybrid crosses. We use data from the F1 generation of our experiment to test the predictions that, relative to pure crosses, hybrid crosses will result in (1) biased sex ratios, (2) lower egg hatching rates, or (3) reduced survival of offspring. We further use data from the backcross generation to test for two forms of hybrid sterility: (1) lower total egg production and (2) lower viable egg production from backcross relative to pure matings.

## 2 Study system

We conduct the first large-scale, multi-generational test for intrinsic isolation in squamate reptiles, a group that includes more than 9,400 species of snakes and lizards (roughly 30% of all terrestrial vertebrates) (Roskov et al. 2015). Very little is known about intrinsic reproductive isolation in this group (but see Arrayago et al. 1996; Corl et al. 2012; Gorman et al. 1971; Olsson et al. 2004; Rykena 1996; Webster 1977). Experimental hybridization studies involving lizards in the genus *Lacerta* report varying degrees intrinsic isolation due to sterility or reduced viability of hybrids (Arrayago et al. 1996; Olsson et al. 2004; Rykena 1996) whereas experiments involving color morphs of side blotched lizards found intrinsic isolation due to reduced production of viable eggs in between-population crosses (Corl et al. 2012).

We use no-choice mating experiments in a common garden environment to investigate the evolution of intrinsic reproductive isolation between two subspecies of an abundant and widely distributed trunk-dwelling anole lizard (*Anolis distichus*) from the island of Hispaniola. This species is particularly well-suited for investigating the evolution of intrinsic isolation during anole speciation for several reasons. First, the two taxa investigated here are distinguished primarily by differences in the color and pattern of the dewlap, but also occupy different environments; the primarily orange-dewlapped *A. d. ignigularis* are endemic to relatively mesic environments in eastern Hispaniola whereas the primarily yellow-dewlapped *A. d. ravitergum* are found only in the more xeric south-central portion of Hispaniola. Although traditionally classified as subspecies due to their tendency to hybridize where they come into contact, more recent studies involving mitochondrial DNA, nuclear microsatellites, and AFLP genome scans all suggest that these taxa are more accurately characterized as a young species pair (Geneva et al. 2015; Glor and Laport 2012; MacGuigan et al. 2017; Ng et al. 2016; Schwartz 1968). These lineages diverged 2.5–5 million years ago and currently meet along a narrow hybrid zone associated with a sharp ecological gradient (Geneva et al. 2015; MacGuigan et al. 2017; Ng and Glor 2011; Ng et al. 2013, 2012, 2016). Absence of intrinsic isolation in the laboratory would suggest that the narrow hybrid zone between these species is maintained entirely by extrinsic mechanisms.

## 3 Methods

### 3.1 Population Abbreviations and Cross Notation

We use abbreviated subspecies names to refer to the two taxa involved in our experiment; ign or *ignigularis* for *Anolis distichus ignigularis* and rav or *ravitergum* for *Anolis distichus ravitergum*. We use male *×* female notation to describe crosses (i.e., rav *×* ign indicates a male rav crossed with a female ign) and male/female notation to describe hybrid individuals (i.e., ign/rav refers to an F1 hybrid with a *A. d. ignigularis* father and a *A. d. ravitergum* mother).

### 3.2 Sampling and husbandry

#### 3.2.1 Sampling of lizards from nature

Between January 1st and 14th, 2011 we sampled sexually mature male and female lizards from two wild populations in the Dominican Republic. These lizards were evenly divided between *A. d. ignigularis* and *A. d. ravitergum*, with each subspecies sampled from a single location. We sampled *A. d. ignigularis* from mesic broadleaf forest along an unpaved road extending north from the western edge of the city of Bańı and the town of El Recodo and running alongside the Rio Bańı (18.3764*^◦^* N, 70.3341*^◦^* W). At this locality, we sampled lizards primarily from trees along the road or river, rocks in the river bed, or two small dilapidated wooden buildings alongside to the road. We sampled *A. d. ravitergum* from a site along the unpaved road extending north from Bańı to the small town of Manaclar that was approximately 6 km (straight line distance) from the *A. d. ignigularis* sampling locality (18.3259*^◦^* N, 70.3464*^◦^* W). Sampling at this locality occurred primarily on buildings and trees associated with a small-scale avocado plantation or from fence posts and trees in the surrounding pasture and xeric scrub forest. We specifically selected these locations because previously acquired mitochondrial, microsatellite and AFLP data found no evidence of individuals with mixed ancestry at these localities which are located at least 2 km from the nearest known hybrid localities (MacGuigan et al. 2017; Ng and Glor 2011). We transported lizards in ventilated 500 mL clear plastic cups (superiorshippingsupplies.com), each of which contained a moist paper towel. These cups were subsequently packed inside a large insulated cooler and shipped to the United States via commercial airliner.

#### 3.2.2 Husbandry of adult lizards

We housed lizards in a climate controlled room maintained as a AAALAC accredited satellite facility of the University of Rochester School of Medicine and Dentistry’s Vivarium. All protocols for lizard husbandry were approved by the University of Rochester’s Animal Care and Use Committee (IACUC protocol number UCAR-2007-030R). The temperature and photoperiod of this facility were set to mimic natural seasonality. During late spring, summer, and early fall we maintained 14 hour light, 10 hour dark light cycle with constant temperature of 85*^◦^*F. From late Fall through early Spring we maintained a 10 hour light, 14 hour dark light cycle with constant temperature of 83*^◦^*F. Humidity of the room was maintained by a steam humidifier set to 50%, although room humidity varied seasonally and humidity tended to be higher within lizard enclosures (Ng et al. 2013).

We lightly misted each enclosure twice daily with distilled water to provide drinking water and to maintain humidity. We fed adult lizards live, two week old crickets obtained from Fluker’s Cricket Farm (Port Allen, LA, USA) three times weekly, twice with crickets dusted with Herptivite® multivitamin supplement (Rep-Cal®, Los Gatos, CA, USA) and once weekly with a 1:1 mix of Herptivite® and Ultrafine Calcium Powder with Vitamin D3 (Rep-Cal®, Los Gatos, CA, USA) (Ascher et al. 2013). We cleaned the interior of each enclosure as well as the dowels and artificial plants as necessary using distilled water. We replaced soiled dowels and plants with enclosure dressings sterilized by soaking them for *>*30 minutes in a 10% bleach solution, followed by soaking and rinsing in distilled water and air drying.

#### 3.2.3 Breeding groups

Subsequent to their arrival at the laboratory facility, males and females were housed separately for 102 days. This period of isolation was necessary because female anoles, like many other lizard species, are capable of sperm storage and may continue to produce eggs for weeks or months after their most recent mating. Estimates of anole sperm storage duration vary widely by species, from less than 10 days to up to 10 months (Passek 2002; Stamps 1974). In over five years of maintaining a large captive colony of *A. distichus*, the longest gap between exposure to a male and production of viable eggs we have observed is 84 days (unpublished data). During this period of sexual isolation following arrival in the lizard facility, *Anolis d. ignigularis* did not produce any eggs. *Anolis d. ravitergum* occasionally produced eggs (approximately 0.026 eggs per female per week) during the first 46 days of the isolation period. Because we continued to house these females in isolation for another 56 days with no further egg production prior to initiating our experiment, it is unlikely that any of the eggs produced during our experiment result from the female’s use of stored sperm.

For each generation of our experiment, we housed lizards in breeding groups comprising one male and two females from the same population. Prior breeding experiments in our facility suggested that this ratio maximizes productivity relative to breeding groups with a single male and female or groups with one male and three or more females per enclosure (Ng et al. 2013). When a breeding individual died during the course of the experiment, we replaced it with another animal from our colony drawn from the same population, but previously housed in isolation from members of the opposite sex. We did this until we exhausted our supply of reserve breeders. For the first generation of our study, we maintained each breeding group in a 11.9 L (29.8 *×* 19.7 *×* 20.3 cm) Lee’s Kritter Keeper™ reptile enclosure (L Schultz Inc., San Marcos, CA). For the second generation, we housed all lizards in custom-made 15.1 L (36 *×* 30 *×* 14 cm) lizard enclosures assembled from clear and white opaque acrylic. This change in housing was necessary to fit all breeding groups and offspring within our animal care facility. The largest straight-line distances within the 11.9 and 15.1 liter cages is 41.1 and 48.9 cm respectively. While the added space might be expected to delay interactions between animals housed in these larger cages *Anolis distichus* can run at least 105 cm per sec (Vanhooydonck et al. 2006). Therefore, interactions between individuals in the 15.1 L cages are delayed by no more than seven hundredths of a second relative to 11.9 L cages (the time to travel the additional 7.8 cm). Each enclosure included 500 mL (*≈*100 grams) of organic potting soil (MiracleGrow™ Scotts Company LLC) as substrate, two 40 cm x 1.25 cm diameter wooden dowel perches, and artificial plants including a total of 8-12 leaves.

All lizard enclosures were arranged into three aisles, two rows, and three shelves. The F1 generation used all three aisles, whereas the backcross generation required only two aisles. Because different enclosure positions in the facility likely resulted in slight differences in enclosure microclimate, we alternated enclosures for each type of breeding group across all aisles, rows, and shelves. We also performed multivariate analyses to examine if the arrangement of enclosures in the vivarium had a significant effect on our measures of reproductive isolation.

#### 3.2.4 Egg laying, incubation, and hatchling husbandry

Throughout the course of our breeding experiments, we provided each breeding group with a egg-laying container made from modified 32oz polypropylene yogurt cups. After cleaning and sterilizing each egg-laying container, we cut 3 cm diameter holes in the center of the lid and on the side of the container, approximately 9 cm from the base. We then filled each egg-laying container up to the side opening with medium grain vermiculite saturated with distilled water, just shy of the point where water dripped from compressed vermiculite. Once a week, we searched for eggs by emptying each egg-laying container and sifting through the vermiculite inside. We also visually inspected the soil and artificial plants in each enclosure for eggs deposited outside the egg-laying container. Upon discovery, we classified eggs as either viable (calcified) or inviable (uncalcified). Prior to returning the egg laying containers to the enclosure, we cleaned them with distilled water and re-moistened the vermiculite.

We individually incubated all viable eggs in ventilated 500 mL clear plastic cups (superiorshippingsupplies.com) containing 130 g of moistened vermiculite (18:11 water to vermiculite by weight). We checked incubating eggs daily for evidence of hatching or egg failure. Eggs that failed during incubation — identified by collapsed egg and/or the presence of mold — were removed from the incubation containers and discarded after noting the date and reason for egg failure. When possible, we dissected these eggs to determine if an embryo was present, and recorded its degree of development.

Immediately following hatching, we uniquely identified hatchlings using a toe clipping scheme modified from Ferner (2007). Hatchlings were then housed in 36 *×* 30 *×* 14 cm enclosures with the same quantity of soil, dowels, and plants as the breeding enclosures. We reared hatchlings at a starting density of up to six lizards per enclosure and in groups of individuals that hatched no more than seven days apart. After one month, enclosures that had less than four animals due to mortality were re-housed with other individuals of similar age to maintain a density of four to six animals per enclosure. No animals were re-housed more than once. We fed hatchlings and checked for hatchling mortality once daily.

Four to six months after hatching, we were able to distinguish male from female hatchlings based on the presence of an extensible dewlap and/or the presence of an enlarged pair of post-anal scales in males. Once this sexing was possible, we re-arranged lizards into new enclosures sorted by sex, with six individuals in female enclosures and four individuals per male enclosure.

### 3.3 Experimental design

To test for evidence of intrinsic reproductive isolation in *A. distichus*, we conducted two generations of no-choice breeding experiments in a controlled laboratory environment. During the first generation of this experiment, we evenly divided adult animals obtained from nature among each of four possible cross types, two pure crosses (♂ *A. d. ignigularis ×* ♀ *A. d. ignigularis* and ♂ *A. d. ravitergum ×* ♀ *A. d. ravitergum*) and each of the reciprocal hybrid crosses (♂ *A. d. ignigularis ×* ♀ *A. d. ravitergum* and ♂ *A. d. ravitergum ×* ♀ *A. d. ignigularis*). During the second generation of the experiment, we conducted backcrosses between pure and F1 hybrid progeny from the first generation cross. We chose to perform backcrosses (in lieu of F2 crosses) to allow for the identification of sex-specific differences in F1 fertility, if present. Finally, to assess egg production of unmated females we established three female-only cages (each with three *A. d. ravitergum*). Female-only cages were limited to *A. d. ravitergum* as we lacked extra *A. d. ignigularis* females.

We quantified egg production based on the number of females in an enclosure at the time of collection (e.g., one egg collected from an enclosure with two females is recorded as 0.5 eggs per female for that collection period). We performed all subsequent statistical analyses on egg counts scaled in this manner. We measured the reproductive output of experimental pairs by calculating the total number of eggs (viable and inviable) produced per female per week in the first (F1) generation of the experiment. We estimated fertilization rates in the F1 generation using the number of viable eggs produced per female per week. We calculated three measures of F1 offspring viability: the portion of viable eggs that produced hatchlings, the survival of hatchlings to 180 days, and the sex ratio of hatchlings. We calculated offspring fertility by calculating the number of total and viable eggs produced per female per week during the backcross generation.

### 3.4 Testing for intrinsic reproductive isolation

To test for significant reproductive isolation between *A. d. ignigularis* and *A. d. ravitergum* we fit generalized linear models (GLM) to each of five response variables predicted to vary in hybrids and their progeny due to intrinsic isolation: (1) total egg production per female per week (F1 and backcross), (2) viable egg production per female per week (F1 and backcross), (3) egg mortality during incubation (F1), (4) hatchling survival to six months (F1), and (5) hatchling sex ratio (F1). In all analyses, we treated breeding group as the unit of replication. Response variables were continuously distributed and modeled as Gaussian random variables, except where noted below.

We used *F* - tests to compare model fit of four hierarchically nested models to each of the five response variables. First, we fit a null model (*M*_0_) designed to test if study design (the arrangement of cages in our lizard facility) produced any unforeseen biases in response variables. This null model (*M*_0_) included three independent variables that describe an enclosure’s location: (1) *X* — front, middle, or back aisle, (2) *Y* — rack on the eastern or western side of the room, and (3) *Z* — bottom, middle, or top shelf. Each of three additional models sequentially adds a single independent variable to the model in the following sequence: model (*M*_1_) adds the subspecies of the female in the enclosure to (*M*_0_), model (*M*_2_) adds the subspecies of the male in an enclosure to (*M*_1_), and (3) a full model (*M*_3_) adds the identity of the cross (the interaction of male and female identity) to (*M*_2_). We used F-tests to compare all models and determine the best fitted model for each response variable. We performed one-way analysis of variance (ANOVA) on the preferred model to determine which (if any) independent variables within that model had a significant effect on the response variable.

Using this approach, we are able to determine if significant variation in response variables were due to factors other than reproductive isolation. For example, significant differences in egg production among crosses could be due to inadvertent bias introduced by study design (enclosure arrangement in the vivarium), or be due lineage specific differences reproductive biology. Bias caused by study design would result in model testing favoring the null model (*M*_0_) and ANOVA identifying a significant effect of one or more descriptions of enclosure arrangement in the vivarium. Biases introduced by lineage-specific differences in reproductive biology would be revealed by model selection favoring either (*M*_1_) or (*M*_2_), and ANOVA favoring either female or male identity (respectively). Finally, evidence of intrinsic reproductive isolation would result in model testing favoring the full model (*M*_3_) and ANOVA identifying cross as a significant factor.

#### 3.4.1 Quantifying degree of intrinsic isolation relative to previously studied examples

To compare the degree of intrinsic reproductive isolation among subspecies of *A. distichus* to that observed in other previously studied species pairs we calculated Coyne and Orr’s (1989) isolation index (*I*) for each generation of the cross as well as a total isolation index across both generations. We do not use the new method for measuring total isolation introduced by Sobel and colleagues (2015) because this method is designed to incorporate both extrinsic and intrinsic isolating barriers.

The strength of intrinsic isolation in the first generation cross (*I*_F1_) is:

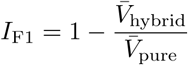

Isolation in the backcross generation (*I*_BC_) is calculated as:

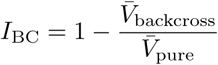

Where *V̅* is the average number of viable eggs per female per week for pure crosses (*V̅*_pure_), hybrid crosses (*V̅*_hybrid_) and backcrosses (*V̅*_backcross_). Earlier acting isolating mechanisms have a proportionally larger effect on fitness (e.g. prezygotic barriers have a stronger effect than postzygotic barrier of equal magnitude) (Ramsey et al. 2003; Schemske 2010). Therefore, the total isolation index is calculated by down-scaling the isolation observed in the backcross as this phenotype reduces the fitness remaining after accounting for the effect observed in the F1.

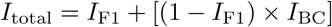

Although there are no statistical tests to assess the significance of this isolation index, we use the values of *I* calculated to contrast anole isolation to that found in other taxa.

### 3.5 Distinguishing among alternative forms of intrinsic reproductive isolation

We used pairwise Mann-Whitney tests applied to the data from our crossing experiment to test the predictions of three possible forms of intrinsic reproductive isolation. We used data from only the first generation of our experiment to test for premating or prezygotic isolation because any reproductive problems during backcrossing could also result from postzygotic reproductive isolation due to partial hybrid sterility. First, we tested premating isolation’s prediction that hybrid crosses would produce significantly fewer eggs overall (including both viable and inviable) than pure crosses during the first generation of our experiment. Second, we tested the prediction that postmating, prezygotic reproductive isolation would result in a reduction in the rate of viable egg production in hybrid crosses relative to pure crosses in the F1 generation. Finally, third, we used data from both generations of our experiment to test for four forms of postzygotic isolation. In the F1 generation we test whether hybrid crosses produced biased sex ratios, lower hatching rates for eggs, and reduced survival of offspring to six months. In the backcross generation we tested if backcross pairings generated lower total egg production and lower viable egg production relative to pure crosses.

## 4 Results

### 4.1 Summary of egg production

#### 4.1.1 First generation F1 cross

In the first generation of our experiment we collected eggs for 37 weeks and ultimately obtained 1702 eggs, of which 1143 were viable and of those, 857 hatched (**Table 1**). This experiment started with 221 animals arranged into 76 breeding groups. Forty-five animals died during the course of the experiment, including 20 (23%) female and nine (20%) male *ignigularis* and 15 (23%) female and one (3%) male *ravitergum*. Breeder mortality was largely uniform throughout the experiment, between 2 and 8 breeders died per month (average 4.4). We replaced these animals with reserve breeders in 16 cages. We did not observe any obvious biases in mortality by cross type with similar mortality between pure (9 and 14 deaths in pure crosses involving *ravitergum* and *ignigularis*, respectively) and hybrid crosses (12 and 10 deaths involving rav *×* ign and ign *×* rav, respectively). We stopped data collection early for three breeding groups (two pure *ignigularis* and one pure *ravitergum*) because we ran out of reserve males. Females in this generation produced an average of 0.45 eggs per week.

**Table 1:** Summary of egg production from both experimental generations.

| Cross Type | ♂ | ♀ | Breeding Groups | Eggs | Total Eggs per ♀ per week (sd) | Viable Eggs per ♀ per week (sd) |
| --- | --- | --- | --- | --- | --- | --- |
| **F1 (first generation)** |  |  |  |  |  |  |
| Pure | ign | ign | 20 | 444 | 0.54 (0.04) | 0.37 (0.14) |
| Pure | rav | rav | 18 | 394 | 0.39 (0.04) | 0.27 (0.09) |
| F1 Hybrid | ign | rav | 20 | 489 | 0.43 (0.03) | 0.18 (0.11) |
| F1 Hybrid | rav | ign | 18 | 375 | 0.43 (0.03) | 0.25 (0.17) |
| **Backcross (second generation)** |  |  |  |  |  |  |
| Backcross | ign | ign/rav | 8 | 139 | 0.42 (0.04) | 0.23 (0.15) |
| Backcross | ign | rav/ign | 6 | 132 | 0.52 (0.06) | 0.32 (0.22) |
| Backcross | rav | ign/rav | 11 | 211 | 0.41 (0.05) | 0.38 (0.15) |
| Backcross | rav | rav/ign | 11 | 168 | 0.37 (0.04) | 0.33 (0.16) |
| Backcross | ign/rav | ign | 2 | 32 | 0.64 (0.07) | 0.54 (0.10) |
| Backcross | rav/ign | ign | 3 | 34 | 0.42 (0.17) | 0.42 (0.30) |
| Backcross | ign/rav | rav | 11 | 161 | 0.32 (0.02) | 0.24 (0.12) |
| Backcross | rav/ign | rav | 11 | 184 | 0.39 (0.03) | 0.27 (0.12) |
| **Summary** |  |  |  |  |  |  |
| Pure crosses |  |  | 38 | 838 | 0.47 (0.17) | 0.33 (0.13) |
| Hybrid crosses |  |  | 38 | 864 | 0.43 (0.13) | 0.21 (0.14) |
| Backcrosses |  |  | 63 | 1061 | 0.40 (0.14) | 0.31 (0.16) |

#### 4.1.2 Second generation backcross

During the second generation of our experiment we collected eggs for 28 weeks and ultimately obtained 1061 eggs, 839 viable eggs, and 679 hatchling lizards from our breeding cages (**Table 1**). This experiment began with 185 individuals arranged into 63 breeding groups. Due to high mortality among pure *ignigularis* hatchlings during the first generation of our experiment, we were not able to investigate as many backcrosses involving F1 hybrids and pure *ignigularis* as backcrosses involving F1 hybrids and pure *ravitergum* (**Table 1**). Throughout the course of this experiment, 20 breeding individuals died, including 17 females and 3 males. Breeder mortality was spread throughout the course of the experiment with 1 to 6 breeders dying per month (average 2.9). We were able to replace seven of these individuals with reserve animals. Mortality rates were comparable across all four types of breeding individuals (pure *ravitergum* −9%, pure *ignigularis* −17%, rav/ign hybrids 14% and ign/rav hybrids 20%). Breeding females in this generation produced an average of 0.40 eggs per week, which was fewer than during the F1 generation but not significantly (*P*_MWU_ = 0.192). The three female-only cages produced exclusively inviable eggs with an average of 0.099 (sd 0.02) eggs per female per week.

### 4.2 Evidence for intrinsic reproductive isolation

#### 4.2.1 GLM hypothesis Testing

We do not find evidence that the arrangement of enclosures significantly influenced any of our measures of reproductive isolation. Although F-tests favored the null model for three response variables (F1 egg mortality, backcross total egg production and backcross viable egg production) our analyses of variance found no evidence that the arrangement of enclosures in the vivarium had a significant effect on any of these measures. In the first generation F1 cross, we found evidence for significant intrinsic reproductive isolation in two response variables: viable egg production and sex ratio (**Table 2**). These measures both favored the complete model (M_3_), and subsequent ANOVA of this model confirmed that both viable egg production and sex ratio were significantly influenced by cross type only. For two additional traits — total egg production and hatchling survival — our analyses significantly favored the M_1_ model, which included population identity of females only. Subsequent ANOVA of the best fit M_1_ model further supported the hypothesis that female identity significantly influenced total egg production and hatchling survival. Pairwise Mann-Whitney analyses (see section 4.3 below), however, suggest that this result is due to variation in total egg production and survival of hatchlings from pure *ignigularis* crosses, and not from reduced egg production or survival of hatchlings from hybrid crosses.

**Table 2:** Summary of GLM analyses. Four nested models were evaluated, each including all independent variables of the previous and adding one additional variable. The null model (M_0_) described study design (e.g. the physical location of breeding groups within the vivarium), M_1_ added the population identity of the females in each cross, M_2_ added male population identity, and M_3_ added the cross as a final factor.

| Response Variable | Preferred Model | d.f. | $F$ | $R^2$ | $P$-value | Significant Factors | $P$-value |
| --- | --- | --- | --- | --- | --- | --- | --- |
| **F1** |  |  |  |  |  |  |  |
| Total eggs | $M_2$ | 67 | 4.88 | 0.193 | 0.031 | female,male | 0.023,0.034 |
| Viable eggs | $M_3$ | 66 | 12.162 | 0.300 | $8.7E^{-4}$ | cross | $2.8E^{-4}$ |
| Egg mortality | $M_0$ | 69 | - | 0.198 | - | - | - |
| 6 mo. Survival | $M_1$ | 68 | 15.363 | 0.257 | $2.1E^{-4}$ | female | $2.4E^{-4}$ |
| Sex Ratio | $M_3$ | 54 | 5.91 | 0.160 | 0.018 | - | - |
| **Backcross** |  |  |  |  |  |  |  |
| Total eggs | $M_0$ | 57 | - | 0.036 | - | - | - |
| Viable eggs | $M_0$ | 57 | - | 0.003 | - | - | - |
| Egg mortality | $M_0$ | 55 | - | 0.073 | - | - | - |
| 6 mo. Survival | $M_1$ | 54 | 4.297 | 0.269 | 0.009 | female | 0.0106 |
| Sex Ratio | $M_0$ | 53 | - | 0.097 | - | - | - |

#### 4.2.2 Measuring isolation

Our GLM model testing of measures of intrinsic reproductive isolation between *Anolis distichus ignigularis* and *A. d. ravitergum* revealed a significant effect of cross on the number of viable eggs and sex ratio (**Table 2, Fig. 1**). Detailed examination of sex ratio data, however, suggests this effect may be an artifact of reduced survival of *A. d. ignigularis* (**Table 1**). As a result, we calculated Coyne and Orr’s (1989) Isolation index for the F1 crosses using only the number of viable eggs produced per female per week resulting in an *I*_F1_ value of 0.343.

**Figure 1:**
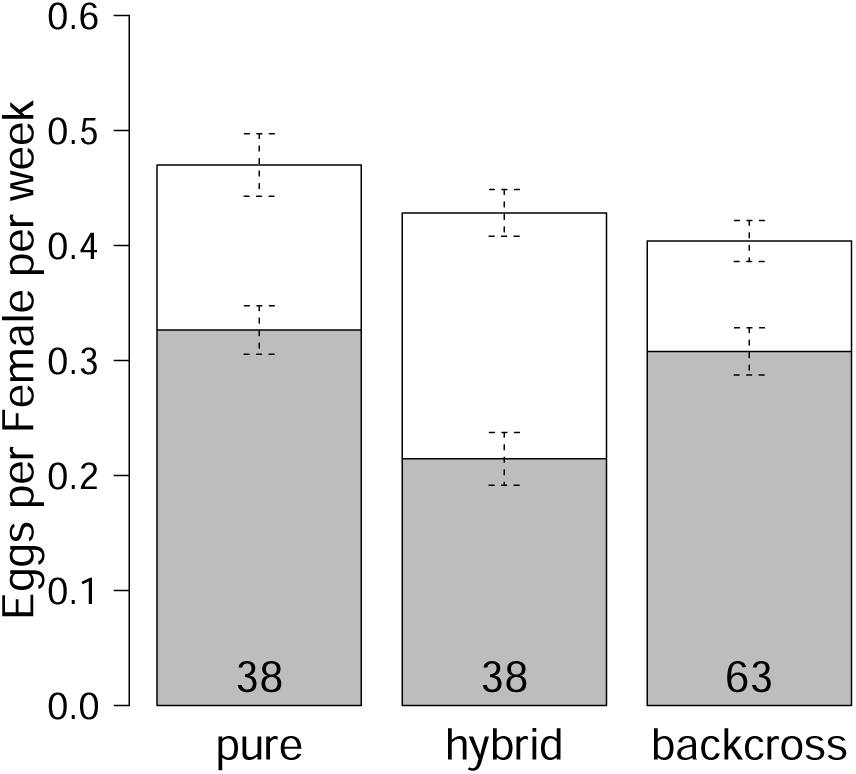
Viable (grey) and inviable (white) egg production from all experimental crosses, grouped by type of cross. The number of replicate crosses used to calculate each mean is listed within each bar. Error bars depict *±*1 standard error of the mean.

Although GLM model testing did not suggest that cross significantly affected backcross viable egg pro-duction (**Table 2**), backcrosses did produce fewer viable eggs than pure crosses (**Table 1**). Therefore, we contrasted pure and backcross viable egg production to estimate the degree to which intrinsic isolation oc-curs in the backcross generation. The isolation index of the backcross alone was substantially smaller than that observed in the F1 generation *I*_BC_ = 0.0570. Combining across both F1 and Backcross generations, we obtain an overall *I*_Total_ value of 0.380.

### 4.3 Characterizing intrinsic isolation

We do not find evidence of premating isolation in the analysis of the total egg production among crosses. Although our GLM analyses suggested female subspecies may contribute to variation in total egg production, the only significant difference in total egg production we observed were between the two pure crosses (*P*_MWU_ = 0.00284). We also find no evidence that hybrid and pure crosses differ in their total egg production (**Table 1**).

We find support for postmating, prezygotic isolation, which predicts a reduction in viable egg production by hybrid crosses. Pure crosses produce more viable eggs than either hybrid cross (**Table 1**). Only the comparison of pure *ignigularis* with the hybrid cross ign *×* rav remains significant after Bonferroni correction for multiple pairwise comparisons. To more directly test for a difference between hybrid and pure crosses we compared pooled pure crosses with pooled hybrid crosses, which reveals a clear, significant difference in viable egg production (*P*_MWU_ = 0.00092), with females assigned to a pure cross producing an average of 4.26 more viable eggs over the course of the 37 week experiment (Fig. 1).

Our analyses find no support for the presence of postzygotic isolation. Incubation success does not differ among crosses and therefore we find no evidence of elevated hybrid inviability during egg incubation (**Table 1**). We observed significantly reduced survival to 6 months of age in the pure *ignigularis* cross (**Table 1**) relative to the ign *×* rav and rav *×* ign hybrid crosses and pure *ravitergum* cross (*P*_MWU_ = 0.00163, 0.00240, 0.00001, respectively). This appears to be a lineage specific effect rather than evidence of hybrid inviability because we found no significant difference in hatchling survival among the pure *ravitergum* and hybrid crosses. Hybrid do not suffer reduced fertility relative to the offspring of pure crosses. Viable and total egg production did not differ significantly between backcrosses and pure populations (**Table S1**).

## 5 Discussion

Our multi-generational crossing experiments involving ecologically divergent subspecies of *Anolis* lizards recover intrinsic reproductive isolation due to a significant decrease in egg viability (Fig. 1). This intrinsic isolation during our first generation hybrid cross results in an isolation index (*I* = 0.343) similar to that observed in the first generation of hybrid crosses between species of *Ficedula* flycatchers (*I* = 0.25 - Price and Bouvier 2002) or between different genera of centrarchid fishes *Ambloplites rupestrus* and *Pomoxis ni-gromaculatus* (*I* = 0.29 - Bolnick and Near 2005). Because the isolation index (*I*) corresponds directly with a decrease in fitness, it is comparable to a selection coefficient, where values as low as 0.01 are expected to generate strong evolutionary responses (reviewed in Charlesworth and Charlesworth 2010). These populations exist at an intermediate stage of reproductive isolation but the strong fitness consequences we observe suggest reproductive isolation may ultimately become complete through the accumulation of further barriers through reinforcing selection against hybridization (Coyne and Orr 2004).

Nevertheless, the fact that we did not recover evidence for significant additional intrinsic reproductive isolation during our second generation backcrossing experiment (*I* = 0.0570, total *I* = 0.398) suggests that F1 hybrids are highly fertile. Moreover, the presence of incomplete intrinsic isolation in crosses between pure parental forms and nearly undetectable intrinsic isolation when F1 hybrids are backcrossed to the pure parental forms suggests that intrinsic barriers alone are likely insufficient to explain the maintenance of ecologically adaptive divergence and limited introgression observed across the narrow hybrid zone between these populations in nature.

Several additional explanations may account for the limited scope of hybridization and introgression in nature. One possibility is hybrid breakdown, where intrinsic isolating barriers have a compounding effect in F2 and later generations. For instance, while the first generation isolation index between *Ficedula* flycatchers is modest, examination of multiple generations reveals virtually complete isolation between these species (Price and Bouvier 2002; Wiley et al. 2009). A second possibility is the existence of extrinsic isolating mechanisms such as: (1) reduced survival of hybrids in nature, potentially due to their possessing an intermediate phenotype that is unfit in the environments occupied by both parental taxa, or (2) reduced mating success of hybrid individuals, potentially due to their possessing a dewlap that is difficult to observe in both parental environments. Finally, while we made every effort to eliminate the effects of extrinsic isolation through our common garden study design, genotype-by-environment interactions cannot be completely eliminated. As a result, it remains possible that some of the signal of reproductive isolation we recover is the result of an extrinsic barrier caused by an interaction with the laboratory environment.

### 5.1 Postmating prezygotic isolation

The fact that the only significant problems we observed in our hybrid crosses involved reduced fertile egg production (and coincident increased infertile egg production) suggests that intrinsic isolation in this pair is due to incomplete post-mating prezygotic isolation rather than premating or postzygotic isolation. We did not recover the significant reduction in total egg production between hybrid and pure crosses predicted by premating isolation (Fig. 1). Incomplete premating isolation could also be responsible for the observed reduction in viable egg production if infrequent mating in hybrid crosses caused insufficient sperm to be available to fertilize a female’s eggs, but two lines of evidence suggest this is not the case. First, hybrid breeding groups in our experiment appear to have been mating throughout the experiment: only one group failed to produce viable eggs and the remaining groups produced a mixture of viable and inviable eggs through the course of the experiment (**Fig. S1**). Second, we do not expect that sperm depletion will be common in mated female anoles because they produce relatively few eggs and have the ability to store large quantities of sperm (Fox 1963; Passek 2002): *Anolis distichus* females in our experiments produced at most 28 eggs, yet females of a congeneric species (*A. carolinensis*) have proven capable of storing up to 189,000 sperm in the storage tubules lining their oviducts (Fox 1963; Passek 2002). Furthermore, exhaustion of stored sperm in hybrid crosses could be an indication of a postmating, prezygotic isolating barrier. If sperm is transferred successfully in between-population crosses, but storage is less efficient than in pure crosses this would represent a postmating, prezygotic barrier resulting from a incompatibility between sperm and the female reproductive tract.

Postzygotic hybrid inviability could also explain the reduced production of viable eggs by hybrid crosses if uncalcified eggs are the result of embryonic failure rather than fertilization failure. Although these alternative are difficult to distinguish with certainty, embryonic failure would have had to occur very early in development because freshly laid calcified eggs always contained visible embryos in early somitogenesis upon dissection whereas uncalcified eggs never contained any visible evidence of embryonic development.

While this study is unable to identify a PMPZ mechanism between *A. d. ignigularis* and *A. d. ravitergum*, the reproductive biology of anoles may suggest an evolutionary process responsible for the formation of PMPZ isolation through sexual conflict. Postmating prezygotic isolation is thought to evolve through sexually antagonistic coevolution driven by sexual conflict within populations (Birkhead and Brillard 2007; Eady 2001; Gavrilets 2000; Parker and Partridge 1998; Swanson and Vacquier 2002). Anoles exhibit at least three forms of sexual conflict: sperm competition, cryptic female choice, and male manipulation of female postcopulatory sexual receptivity (Calsbeek and Bonneaud 2008; Calsbeek et al. 2007; Crews 1973; Johnson 2007; Passek 2002).

The last of these is of particular interest because a similar phenomenon it is thought to lead to PMPZ isolation in *Drosophila* (Chapman 2001; Knowles and Markow 2001; Sweigart 2010; Wigby and Chapman 2005). In *A. carolinensis*, females paired with males whose hemipenes have been removed are sexually receptive within 3-5 minutes while those mated to intact males refuse further mating for at least 24 hours, potentially increasing the likelihood of a male siring that female’s offspring (Crews 1973). In some fly species, male reproductive alleles have evolved that increase the length of time after mating that a female will refuse mating, thereby increasing the portion of that female’s offspring a male sires, but as a byproduct shortens the females lifespan, negatively affecting that female’s fitness (Rice 2000; Stockley 1997). In response, female beneficial alleles have arisen that counteract the effects induced by males (Knowles and Markow 2001). Such conflict can result in a reproductive arms race featuring alternating evolution of male and female beneficial alleles (Chapman et al. 2003; Parker and Partridge 1998). Postmating, prezygotic isolation arises when animals from populations experiencing such arms races hybridize with animals from populations that have not experienced the same sexual co-evolution (Birkhead and Brillard 2007; Eady 2001; Parker and Partridge 1998). If male manipulation of female postcopulatory receptivity has resulted in a sexual-conflict-driven, co-evolutionary arms race in anoles, it could represent a mechanism for the evolution of the PMPZ isolation we observe.

### 5.2 Conclusions

We find evidence of significant intrinsic reproductive isolation between recently diverged and ecologically distinct lizard populations. The strength of the intrinsic isolation we observe is similar to that observed between well-studied populations typically regarding as good species, such as *Ficedula* flycatchers (Price and Bouvier 2002). Our results suggest that the impact of intrinsic reproductive isolation between young, ecologically diverged populations should not be discounted in favor of purely extrinsic explanations. Our findings also suggest that intrinsic isolation we observe results from postmating, prezygotic isolation. This study is the largest reciprocal hybrid cross to directly test for evidence of intrinsic isolation in reptiles, and among the first to use multiple experimental generations. Our findings, and the potential for comparative analyses across the *Anolis* adaptive radiation, suggest that further work in this group may provide valuable insights into the process of speciation.

## 6 Acknowledgments

Collection permissions were granted by the Ministry of Environment and Natural Resources of the Dominican Republic. Daniel Scantlebury and Miguel Landestoy helped with field collection. This work was made possible thanks to the many people who provided countless hours of lizard room assistance, particularly Juli-enne Ng, Dan Scantlebury, Shea Lambert, Shannon Keating, Benjamin Desch, Daniel MacGuigan, Sabina Noll, Audrey Kelly, Frank Chang, and Ryane Logsdon. Dan Scantlebury designed the custom lizard enclosures used in this experiment. Joel McGlothlin, members of the Glor and Losos labs, and two anonymous reviewers provided valuable feedback on earlier versions of this manuscript. Statistical support was provided by data science specialist Simo Goshev, at the Institute for Quantitative Social Science, Harvard University. Funding was provided by NSF DEB #0920892 (REG), NSF DEB #1457774 (REG), NSF DEB #1500761 (REG and AJG), NSF DEB #1927194 (AJG), and a Sproull University Fellowship from the University of Rochester (AJG).

## Supplementary Material

**Table S1:**
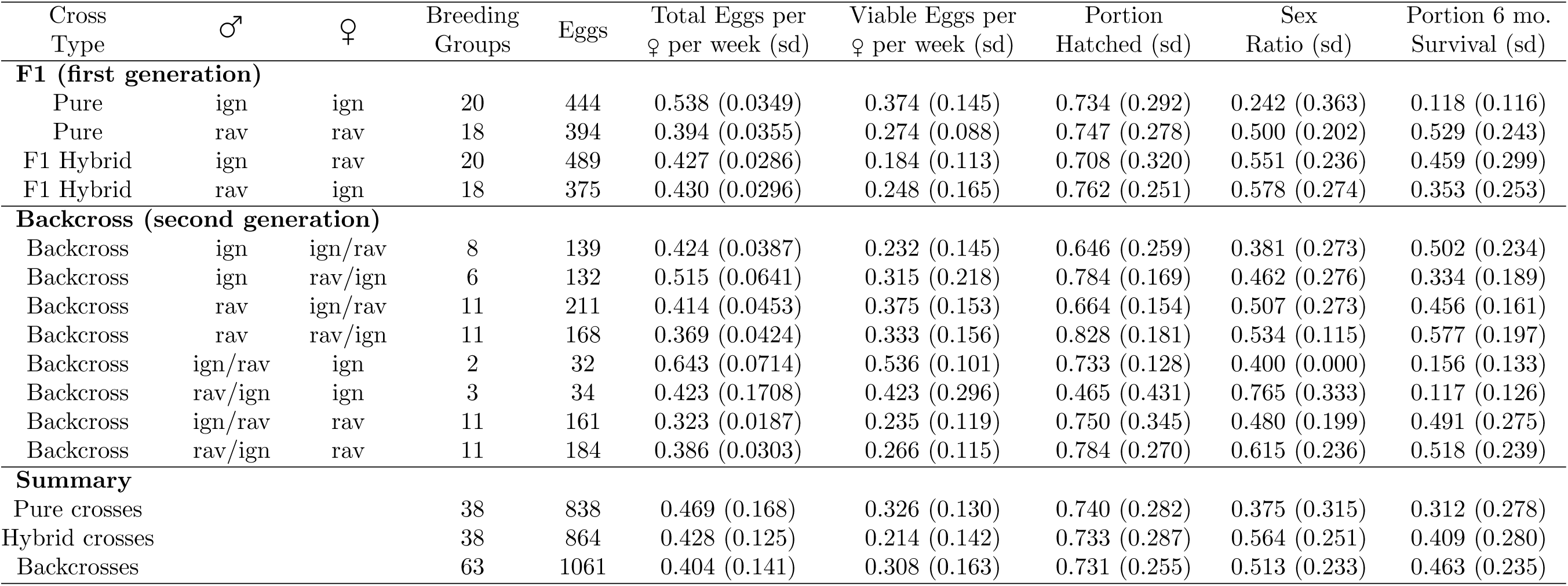
Summary of all phenotypes measured both experimental generations.

| Cross Type | ♂ | ♀ | Breeding Groups | Eggs | Total Eggs per ♀ per week (sd) | Viable Eggs per ♀ per week (sd) | Portion Hatched (sd) | Sex Ratio (sd) | Portion 6 mo. Survival (sd) |
| --- | --- | --- | --- | --- | --- | --- | --- | --- | --- |
| **F1 (first generation)** |  |  |  |  |  |  |  |  |  |
| Pure | ign | ign | 20 | 444 | 0.538 (0.0349) | 0.374 (0.145) | 0.734 (0.292) | 0.242 (0.363) | 0.118 (0.116) |
| Pure | rav | rav | 18 | 394 | 0.394 (0.0355) | 0.274 (0.088) | 0.747 (0.278) | 0.500 (0.202) | 0.529 (0.243) |
| F1 Hybrid | ign | rav | 20 | 489 | 0.427 (0.0286) | 0.184 (0.113) | 0.708 (0.320) | 0.551 (0.236) | 0.459 (0.299) |
| F1 Hybrid | rav | ign | 18 | 375 | 0.430 (0.0296) | 0.248 (0.165) | 0.762 (0.251) | 0.578 (0.274) | 0.353 (0.253) |
| **Backcross (second generation)** |  |  |  |  |  |  |  |  |  |
| Backcross | ign | ign/rav | 8 | 139 | 0.424 (0.0387) | 0.232 (0.145) | 0.646 (0.259) | 0.381 (0.273) | 0.502 (0.234) |
| Backcross | ign | rav/ign | 6 | 132 | 0.515 (0.0641) | 0.315 (0.218) | 0.784 (0.169) | 0.462 (0.276) | 0.334 (0.189) |
| Backcross | rav | ign/rav | 11 | 211 | 0.414 (0.0453) | 0.375 (0.153) | 0.664 (0.154) | 0.507 (0.273) | 0.456 (0.161) |
| Backcross | rav | rav/ign | 11 | 168 | 0.369 (0.0424) | 0.333 (0.156) | 0.828 (0.181) | 0.534 (0.115) | 0.577 (0.197) |
| Backcross | ign/rav | ign | 2 | 32 | 0.643 (0.0714) | 0.536 (0.101) | 0.733 (0.128) | 0.400 (0.000) | 0.156 (0.133) |
| Backcross | rav/ign | ign | 3 | 34 | 0.423 (0.1708) | 0.423 (0.296) | 0.465 (0.431) | 0.765 (0.333) | 0.117 (0.126) |
| Backcross | ign/rav | rav | 11 | 161 | 0.323 (0.0187) | 0.235 (0.119) | 0.750 (0.345) | 0.480 (0.199) | 0.491 (0.275) |
| Backcross | rav/ign | rav | 11 | 184 | 0.386 (0.0303) | 0.266 (0.115) | 0.784 (0.270) | 0.615 (0.236) | 0.518 (0.239) |
| **Summary** |  |  |  |  |  |  |  |  |  |
| Pure crosses |  |  | 38 | 838 | 0.469 (0.168) | 0.326 (0.130) | 0.740 (0.282) | 0.375 (0.315) | 0.312 (0.278) |
| Hybrid crosses |  |  | 38 | 864 | 0.428 (0.125) | 0.214 (0.142) | 0.733 (0.287) | 0.564 (0.251) | 0.409 (0.280) |
| Backcrosses |  |  | 63 | 1061 | 0.404 (0.141) | 0.308 (0.163) | 0.731 (0.255) | 0.513 (0.233) | 0.463 (0.235) |

**Table S2:** Summary of sex ratio for each cross type. P-values calculated via binomial test, testing departure from equal sex ratios.

| Cross Type | ♂ | ♀ | Males | Females | P-value |
| --- | --- | --- | --- | --- | --- |
| **F1 (first generation)** |  |  |  |  |  |
| Pure | ign | ign | 24 | 7 | 0.003 |
| Pure | rav | rav | 96 | 82 | 0.330 |
| F1 Hybrid | ign | rav | 52 | 57 | 0.702 |
| F1 Hybrid | rav | ign | 39 | 46 | 0.515 |
| **Backcross (second generation)** |  |  |  |  |  |
| Backcross | ign | ign/rav | 14 | 16 | 0.856 |
| Backcross | ign | rav/ign | 12 | 16 | 0.572 |
| Backcross | ign/rav | ign | 2 | 1 | 1.000 |
| Backcross | ign/rav | rav | 31 | 36 | 0.625 |
| Backcross | rav/ign | ign | 3 | 5 | 0.727 |
| Backcross | rav/ign | rav | 29 | 35 | 0.532 |
| Backcross | rav | ign/rav | 40 | 49 | 0.397 |
| Backcross | rav | rav/ign | 40 | 50 | 0.343 |

**Figure S1:**
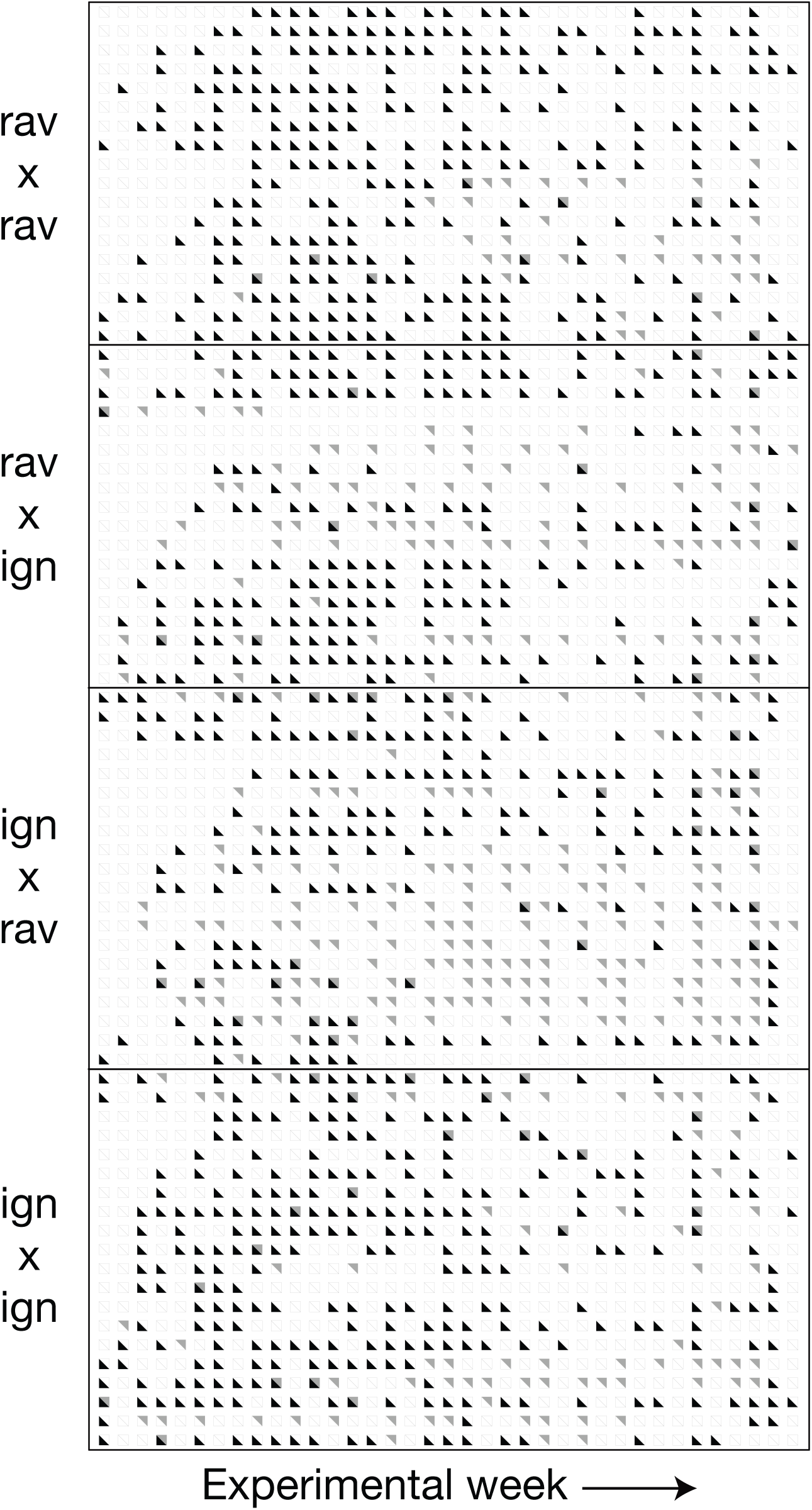
The production of viable and inviable eggs for each cage throughout the F1 experiment. Each row represents the egg output of an experimental cage, grouped by cross type. Columns correspond to each of the 37 weeks in the F1 experiment. Grey triangles represent the production of one or more inviable eggs. Black triangles indicate the production of one or more viable eggs.

## References

Andrews, R. and A. S. Rand, 1974. Reproductive effort in anoline lizards. Ecology 55:1317–1327.

Arrayago, M. J., A. Bea, and B. Heulin, 1996. Hybridization experiment between oviparous and viviparous strains of *Lacerta vivipara*: A new insight into the evolution of viviparity in reptiles. Herpetologica 52:333–342.

Ascher, J. M., A. J. Geneva, J. Ng, J. D. Wyatt, and R. E. Glor, 2013. Phylogenetic analyses of novel squamate adenovirus sequences in wild-caught *Anolis* lizards. PloS one 8:e60977.

Barbash, D. A., D. F. Siino, A. M. Tarone, and J. Roote, 2003. A rapidly evolving myb-related protein causes species isolation in *Drosophila*. Proceedings of the National Academy of Sciences 100:5302–5307.

Birkhead, T. R. and J.-P. Brillard, 2007. Reproductive isolation in birds: postcopulatory prezygotic barriers. Trends In Ecology and Evolution 22:266–272.

Birkhead, T. R., B. C. Sheldon, and F. Fletcher, 1994. A comparative study of sperm–egg interactions in birds. Journal of Reproduction and Fertility 101:353–361.

Bolnick, D. I. and T. J. Near, 2005. Tempo of hybrid inviability in centrarchid fishes (Teleostei: Centrarchidae). Evolution 59:1754–1767.

Brideau, N. J., H. A. Flores, J. Wang, S. Maheshwari, X. Wang, and D. A. Barbash, 2006. Two Dobzhansky-Muller genes interact to cause hybrid lethality in *Drosophila*. Science 314:1292–1295.

Calsbeek, R. and C. Bonneaud, 2008. Postcopulatory fertilization bias as a form of cryptic sexual selection. Evolution 62:1137–1148.

Calsbeek, R., C. Bonneaud, S. Prabhu, N. Manoukis, and T. B. Smith, 2007. Multiple paternity and sperm storage lead to increased genetic diversity in *Anolis* lizards. Evolutionary Ecology Research 9:495–503.

Camargo, A. and J. J. Sites, 2013. Species delimitation: A decade after the renaissance. in I. Pavlinov, ed. The Species Problem - Ongoing Issues. InTech.

Chapman, T., 2001. Seminal fluid-mediated fitness traits in *Drosophila*. Heredity 87:511–521.

Chapman, T., G. Arnqvist, J. Bangham, and L. Rowe, 2003. Sexual conflict. Trends in Ecology and Evolution 18:41–47.

Charlesworth, B. and D. Charlesworth, 2010. Elements of Evolutionary Genetics. Roberts and Company Publishers.

Corl, A., L. T. Lancaster, and B. Sinervo, 2012. Rapid formation of reproductive isolation between two populations of side-blotched lizards, *Uta stansburiana*. Copeia 2012:593–602.

Coyne, J. and H. A. Orr, 2004. Speciation. Sinauer, Sunderland, MA.

Coyne, J. A. and H. A. Orr, 1989. Two rules of speciation. Pp. 180–207, in D. Otte and J. Endler, eds. Speciation and its Consequences. Sunderland, MA, Sinauer.

Crews, D., 1973. Coition-induced inhibition of sexual receptivity in female lizards (*Anolis carolinensis*). Physiology and Behavior 11:463–468.

Dobzhansky, T., 1935. A critique of the species concept in biology. Philosophy of Science Pp. 344–355.

Eady, P. E., 2001. Postcopulatory, prezygotic reproductive isolation. Journal of Zoology 253:47–52.

Ferner, J. W., 2007. A review of marking techniques for amphibians and reptiles. Herpetological circular. Society for the Study of Amphibians and Reptiles.

Fishman, L. and J. H. Willis, 2001. Evidence for Dobzhansky-Muller incompatibilities contributing to the sterility of hybrids between *Mimulus guttatus* and *M. nasutus*. Evolution 55:1932–1942.

Fitzpatrick, B., 2008. Dobzhansky-Muller model of hybrid dysfunction supported by poor burst-speed performance in hybrid tiger salamanders. Journal of evolutionary biology 21:342–351.

Fox, W., 1963. Special tubules for sperm storage in female lizards. Nature 198:500–501.

Gavrilets, S., 2000. Rapid evolution of reproductive barriers driven by sexual conflict. Nature 403:886–889.

Geneva, A., J. Hilton, S. Noll, and R. E. Glor, 2015. Multilocus phylogenetic analyses of Hispaniolan and Bahamian trunk anoles (*distichus* species group). Molecular Phylogenetics and Evolution 87:105–117.

Glor, R. E. and R. G. Laport, 2012. Are subspecies of *Anolis* lizards that differ in dewlap color and pattern also genetically distinct? a mitochondrial analysis. Molecular Phylogenetics and Evolution 64:255–260.

Gorman, G., P. Licht, H. Dessauer, and J. Boos, 1971. Reproductive failure among the hybridizing *Anolis* lizards of Trinidad. Systematic Zoology 20:1–18.

Grant, P. R. and B. R. Grant, 1997. Genetics and the origin of bird species. Proceedings Of The National Academy Of Sciences USA 94:7768–7775.

Hendry, A. P., P. Nosil, and L. H. Rieseberg, 2007. The speed of ecological speciation. Functional Ecology 21:455–464.

Johnson, M. A., 2007. Behavioral ecology of Caribbean *Anolis* lizards: A comparative approach. Ph.D. thesis, Washington University.

Kamath, A. and J. Losos, 2017. Integrating movement behavior with sexual selection in the lizard *Anolis sagrei*. bioRxiv 164863.

Knowles, L. L. and T. A. Markow, 2001. Sexually antagonistic coevolution of a postmating-prezygotic reproductive character in desert *Drosophila*. Proceedings of the National Academy of Sciences 98:8692– 8696.

Larson, E. L., G. L. Hume, J. A. Andŕes, and R. G. Harrison, 2012. Post-mating prezygotic barriers to gene exchange between hybridizing field crickets. Journal of Evolutionary Biology 25:174–186.

Leal, M. and L. J. Fleishman, 2004. Differences in visual signal design and detectability between allopatric populations of *Anolis* lizards. The American Naturalist 163:26–39.

Littlejohn, M., 1965. Premating isolation in the *Hyla ewingi* complex (Anura: Hylidae). Evolution 19:234– 243.

Losos, J., 2009. Lizards in an Evolutionary Tree: Ecology and Adaptive Radiation of Anoles. Organisms and Environments. University of California Press.

Losos, J. B., 1985. An experimental demonstration of the species-recognition role of *Anolis* dewlap color. Copeia Pp. 905–910.

MacGuigan, D. J., A. J. Geneva, and R. E. Glor, 2017. A genomic assessment of species boundaries and hybridization in a group of highly polymorphic anoles (*distichus* species complex). Ecology and Evolution 7:3657–3671.

Mayr, E., 1963. Animal species and evolution. Harvard University Press, Cambridge, Massachusetts.

Mayr, E., 1992. A local flora and the biological species concept. American Journal of Botany 79:222–238.

Moyle, L. C. and T. Nakazato, 2009. Complex epistasis for Dobzhansky–Muller hybrid incompatibility in *Solanum*. Genetics 181:347–351.

Ng, J., A. Geneva, S. Noll, and R. Glor, 2017. Signals and speciation: *Anolis* dewlap color as a reproductive barrier. Journal of Herpetology 51:437–447.

Ng, J. and R. E. Glor, 2011. Genetic differentiation among populations of a Hispaniolan trunk anole that exhibit geographical variation in dewlap colour. Molecular Ecology 20:4302–4317.

Ng, J., A. L. Kelly, D. J. MacGuigan, and R. E. Glor, 2013. The role of heritable and dietary factors in the sexual signal of a Hispaniolan *Anolis* lizard, *Anolis distichus*. Journal of Heredity 104:862–873.

Ng, J., E. L. Landeen, R. M. Logsdon, and R. E. Glor, 2012. Correlation between *Anolis* lizard dewlap phe-notype and evironmental variation indicates adaptive divergence of a signal important to sexual selection and species recognition. Evolution 67:573–582.

Ng, J., A. G. Ossip-Klein, and R. E. Glor, 2016. Adaptive signal coloration maintained in the face of gene flow in a Hispaniolan *Anolis* Lizard. BMC Evolutionary Biology 16:1–12.

Nosil, P., 2012. Ecological speciation. Oxford University Press.

Nosil, P., T. H. Vines, and D. J. Funk, 2005. Reproductive isolation caused by natural selection against immigrants from divergent habitats. Evolution 59:705–719.

Olsson, M., A. Gullberg, and H. Tegelströ, 1994. Sperm competition in the sand lizard, *Lacerta agilis*. Animal Behaviour 48:193–200.

Olsson, M. and R. Shine, 1997. Advantages of multiple matings to females: a test of the infertility hypothesis using lizards. Evolution 51:1684–1688.

Olsson, M., B. Ujvari, T. Madsen, T. Uller, and E. Wapstra, 2004. Haldane rules: costs of outbreeding at production of daughters in sand lizards. Ecology Letters 7:924–928.

Orr, H. A., 1995. The population genetics of speciation: the evolution of hybrid incompatibilities. Genetics 139:1805–1813.

Orr, H. A. and M. Turelli, 2001. The evolution of postzygotic isolation: Accumulating Dobzhansky-Muller incompatibilities. Evolution 55:1085.

Parker, G. A. and L. Partridge, 1998. Sexual conflict and speciation. Philosophical Transactions of the Royal Society of London B: Biological Sciences 353:261–274.

Passek, K. M., 2002. Extra-pair paternity within the female-defense polygyny of the lizard, *Anolis carolinensis*: Evidence of alternative mating strategies. Ph.D. thesis, Virginia Polytechnic Institute and State University.

Presgraves, D. C., 2010. The molecular evolutionary basis of species formation. Nature Reviews Genetics 11:175–180.

Price, C. S., C. H. Kim, C. J. Gronlund, and J. A. Coyne, 2001. Cryptic reproductive isolation in the *Drosophila simulans* species complex. Evolution 55:81–92.

Price, T. D. and M. M. Bouvier, 2002. The evolution of F1 postzygotic incompatibilities in birds. Evolution 56:2083–2089.

Ramsey, J., H. D. Bradshaw, and D. W. Schemske, 2003. Components of reproductive isolation between the monkeyflowers *Mimulus lewisii* and *M. cardinalis* (Phrymaceae). Evolution 57:1520–1534.

Rice, W. R., 2000. Dangerous liaisons. Proceedings of the National Academy of Sciences 97:12953–12955.

Roskov, Y., L. Abucay, T. Orrell, D. Nicolson, T. Kunze, A. Culham, N. Bailly, P. Kirk, T. Bourgoin, R. E. DeWalt, W. Decock, and A. De Wever (eds.) 2015. Species 2000 and ITIS Catalogue of Life. Species 2000: Naturalis, Leiden, the Netherlands. URL www.catalogueoflife.org.

Rykena, S., 1996. Experimental interspecific hybridization in the genus *Lacerta*. Israel Journal of Zoology 42:171–184.

Schemske, D. W., 2010. Adaptation and the origin of species. The American Naturalist 176:S4–S25.

Schluter, D., 2009. Evidence for ecological speciation and its alternative. Science 323:737.

Schuluter, D., 2000. The ecology of adaptive radiation. Oxford University Press, Oxford.

Schwartz, A., 1968. Geographic variation in *Anolis distichus* Cope (Lacertilia, Iguanidae) in the Bahama Islands and Hispaniola. Bulletin of The Museum of Comparative Zoology 137:255–309.

Scopece, G., A. Musacchio, A. Widmer, and S. Cozzolino, 2007. Patterns of reproductive isolation in mediterranean deceptive orchids. Evolution 61:2623–2642.

Seehausen, O., G. Takimoto, D. Roy, and J. Jokela, 2008. Speciation reversal and biodiversity dynamics with hybridization in changing environments. Molecular Ecology 17:30–44.

Seehausen, O., J. vanAlphen, and F. Witte, 1997. Cichlid fish diversity threatened by eutrophication that curbs sexual selection. Science 277:1808–1811.

Sinervo, B. and P. Licht, 1991. Hormonal and physiological control of clutch size, egg size, and egg shape in side-blotched lizards (*Uta stansburiana*: constraints on the evolution of lizard life histories. Journal of Experimental Zoology 257:252–264.

Sites, J. W. and J. C. Marshall, 2003. Delimiting species: a renaissance issue in systematic biology. Trends in Ecology & Evolution 18:462–470.

Smadja, C. and R. Butlin, 2009. On the scent of speciation: The chemosensory system and its role in premating isolation. Heredity 102:77–97.

Sobel, J. M., G. F. Chen, L. R. Watt, and D. W. Schemske, 2010. The Biology of Speciation. Evolution 64:295–315.

Sobel, J. M. and M. A. Streisfeld, 2015. Strong premating reproductive isolation drives incipient speciation in *Mimulus aurantiacus*. Evolution 69:447–461.

Stamps, J. A., 1974. Courtship patterns, estrus periods and reproductive condition in a lizard, *Anolis aeneus*. Physiology and Behavior 14:531–535.

Stockley, P., 1997. Sexual conflict resulting from adaptations to sperm competition. Trends in Ecology and Evolution 12:154–159.

Swanson, W. J. and V. D. Vacquier, 2002. The rapid evolution of reproductive proteins. Nature Reviews Genetics 3:137–144.

Sweigart, A. L., 2010. The genetics of postmating, prezygotic reproductive isolation between *Drosophila virilis* and *D. americana*. Genetics 184:401–410.

Turelli, M., N. H. Barton, and J. A. Coyne, 2001. Theory and speciation. Trends in Ecology and Evolution 16:330–343.

Vacquier, V. D. and Y.-H. Lee, 1993. Abalone sperm lysin: unusual mode of evolution of a gamete recognition protein. Zygote 1:181–196.

Vanhooydonck, B., A. Herrel, R. Van Damme, and D. J. Irschick, 2006. The quick and the fast: The evolution of acceleration capacity in *Anolis* lizards. Evolution 60:2137–12.

Wade, M. J., H. Patterson, N. W. Chang, and N. A. Johnson, 1994. Postcopulatory, prezygotic isolation in flour beetles. Heredity 72:163–167.

Webster, T. P., 1974. Report. Pp. 1–4, in E. E. Williams, ed. The second Anolis newsletter. Cambridge, MA.

Webster, T. P., 1977. Hybridization of Hispaniolan lizards in the Anolis distichus species group. Pp. 166–173, in E. E. Williams, ed. The third Anolis newsletter. Cambridge, MA.

Whittemore, A. T., 1993. Species concepts: a reply to Ernst Mayr. Taxon 42:573–583.

Wigby, S. and T. Chapman, 2005. Sex peptide causes mating costs in female *Drosophila melanogaster*. Current Biology 15:316–321.

Wiley, C., A. Qvarnström, G. Andersson, T. Borge, and G.-P. Saetre, 2009. Postzygotic isolation over multiple generations of hybrid descendents in a natural hybrid zone: How well do single-generation estimates reflect reproductive isolation? Evolution 63:1731–1739.

Williams, E. E., 1972. The origins of faunas. Evolution of lizard congeners in a complex island fauna: a trial analysis. Evolutionary Biology 6:47–89.

Williams, E. E. and A. S. Rand, 1977. Species recognition, dewlap function and faunal size. American Zoologist 17:261–270.

